# Convergent patterns of gene expression and protein evolution associated with adaptation to desert environments in rodents

**DOI:** 10.1101/2021.09.10.459863

**Authors:** Noëlle K. J. Bittner, Katya L. Mack, Michael W. Nachman

**Author notes:** Corresponding Author: Michael Nachman, Department of Integrative Biology and Museum of Vertebrate Zoology, 3101 Valley Life Sciences Building, University of California Berkeley, Berkeley, CA 94720.

## Abstract

Desert specialization has arisen multiple times across rodents and is often associated with a suite of convergent phenotypes, including modification of the kidneys to mitigate water loss. However, the extent to which phenotypic convergence in desert rodents is mirrored at the molecular level is unknown. Here, we sequenced kidney mRNA and assembled transcriptomes for three pairs of rodent species to search for convergence in gene expression and amino acid sequence associated with adaptation to deserts. We conducted phylogenetically-independent comparisons between a desert specialist and a non-desert relative in three families representing ∼70 million years of evolution. Overall, patterns of gene expression faithfully recapitulated the phylogeny of these six taxa. However, we found that 8.6% of all genes showed convergent patterns of expression evolution between desert and non-desert taxa, a proportion that is much higher than expected by chance. In addition to these convergent changes, we observed many species-pair specific changes in gene expression indicating that different instances of adaptation to deserts include a combination of unique and shared changes. Patterns of protein evolution revealed a small number of genes showing evidence of positive selection, the majority of which did not show convergent changes in gene expression. Overall, our results suggest convergent changes in gene regulation play a primary role in the complex trait of desert adaptation in rodents.

## Introduction

The repeatability of adaptive evolution at the molecular level remains an open question. In situations where the mutational target is small and constraints exist due to epistasis or pleiotropy, the molecular paths available to adaptation may be highly limited (Weinreich et al. 2006; Karageorgi et al. 2019). Indeed, there are a number of excellent examples of convergent molecular evolution underlying simple traits (e.g. Stewart and Wilson 1987; Mundy 2005; Zhen et al. 2012). For highly polygenic traits, however, convergence may be less expected simply because the mutational target is large and multiple paths may be available on which selection can act. Nonetheless, several studies have found evidence for convergence at the molecular level even for complex traits (e.g. Marcovitz et al. 2019; Sackton et al. 2019).

Convergent phenotypic evolution may be due to changes in gene regulation, to changes in protein structure, or both, yet these processes are rarely studied together in the context of complex adaptive traits (but see Hao et al. 2019). There is evidence that gene expression divergence and amino acid sequence divergence are correlated between paralogs following gene duplications (Gu et al. 2002; Makova and Li 2003), and more generally that rates of gene expression and rates of protein evolution are coupled in some lineages (e.g. Nuzhdin et al. 2004; Lemos et al. 2005). These observations raise the possibility that changes in both gene expression and protein sequence may contribute to the repeated evolution of complex adaptive traits.

Adaptation to desert environments in rodents provides an opportunity to study repeated evolution in both gene expression and protein sequence in a complex trait. Desert ecosystems present the challenge of extreme aridity and low or seasonally absent water, yet multiple lineages of rodents have independently evolved the ability to survive in these unusually harsh environments (reviewed in Degen 1997). Rodents have solved these challenges in myriad ways, including dietary specialization on plants that are relatively high in water content or modifications to reduce evaporative water loss (Schmidt-Nielsen and Schmidt-Nielsen 1952; Schmidt-Nielsen 1964; Degen 1997). However, a common feature of most desert rodents is a modified kidney capable of producing highly concentrated urine (MacMillen and Lee 1967; Beuchat 1990; Al-kahtani et al. 2004; Donald and Pannabecker 2015). Final excreted urine concentration depends on the development and maintenance of a corticomedullary osmotic gradient within the kidney. Studies have shown that many aspects of kidney morphology and physiology have been modified in different lineages to produce hyper-concentrated urine (Bankir and de Rouffignac 1985; Donald and Pannabecker 2015).

The genetic basis of desert adaptation has been studied independently in a handful of species and individual genes and pathways which may underlie this adaptive phenotype have been identified (Rocha et al. 2021). Here, we leverage three phylogenetically independent lineages of rodents that have all converged on a common phenotype, ultra-high urine concentration associated with desert living, to identify shared molecular changes associated with habitat type. We compared kidney gene expression and protein sequence divergence between a desert and a non-desert species in each of three pairs of phylogenetically independent comparisons representing transitions to desert living in three different rodent families (Heteromyidae, Dipodidae, and Muridae). Desert species were chosen based on their high urine concentration, a proxy for increased osmoregulatory capacity (Figure 1). Within Muridae, we compared the Australian Spinifex Hopping Mouse, *Notomys alexis*, the mammal with the highest known urine concentration and well studied for its modifications to desert life (MacMillen and Lee 1967; Macmillen and Lee 1969; Baudinette 1972; Donald et al. 2012), to the house mouse (*Mus musculus*), a widespread generalist. Within Dipodidae, we compared the desert-dwelling Lesser Egyptian Jerboa, *Jaculus jaculus*, previously studied for its kidney modifications associated with granivorous desert living (Schmidt-Nielsen and Schmidt-Nielsen 1952; Khalil and Tawfic 1963), to the Western Jumping Mouse, *Zapus princeps*, a North American species found in riparian environments. Within Heteromyidae, we compared the Rock Pocket Mouse, *Chaetodipus intermedius* (Bradley et al. 1975; Altschuler et al. 1979), native to the North American Sonoran desert, to the Desmarest’s spiny pocket mouse, *Heteromys desmarestianus*, a neotropical species found in mesic areas that cannot survive without free water (Fleming 1977).

**Figure 1.**
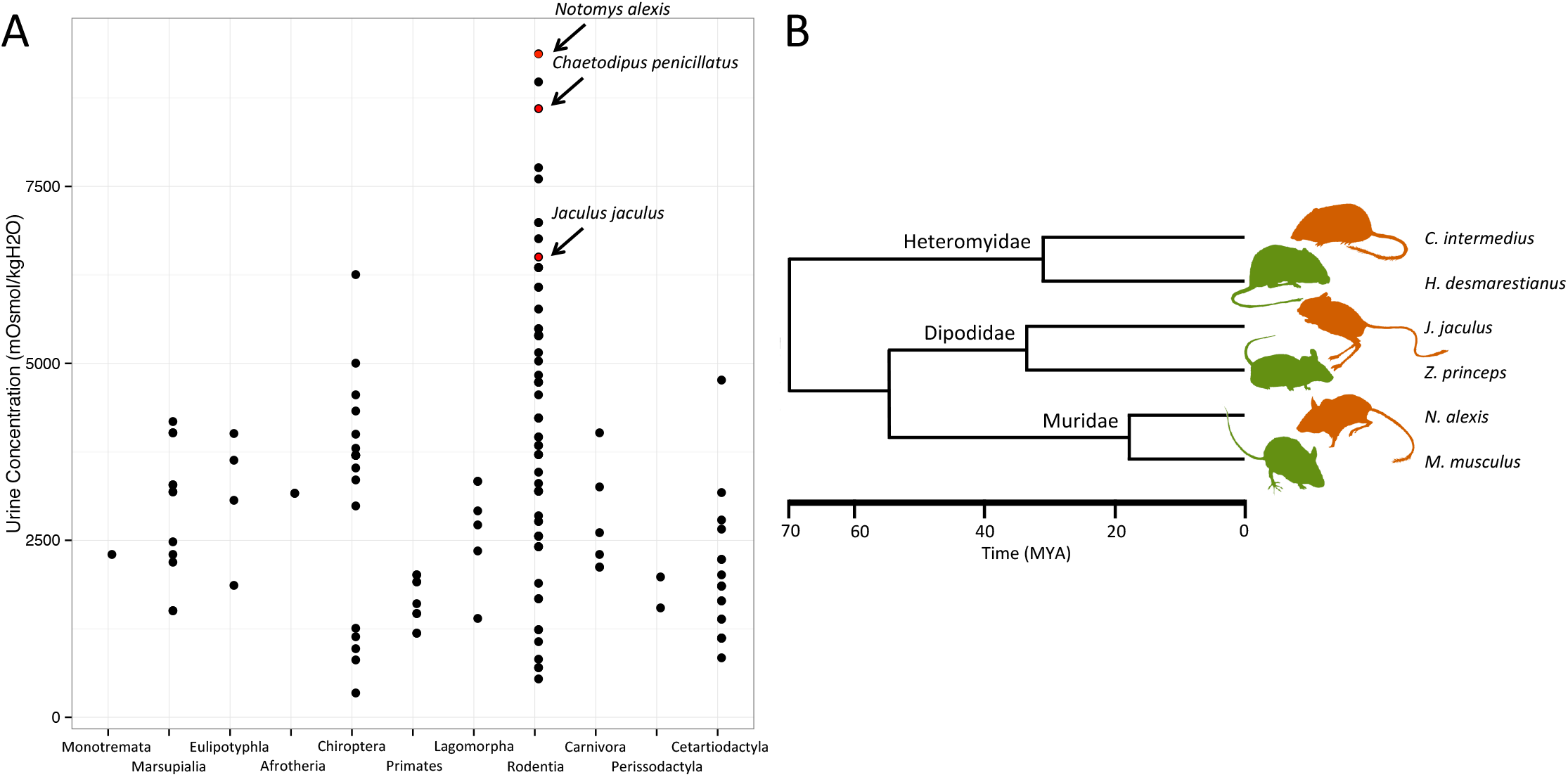
A. Estimates of urine concentration from across Mammalia from Beuchat (1990). Notably, Rodentia has representatives with the highest urine concentrations recorded in mammals. The three desert specialists in this study have among the highest urine concentrations measured in rodents. Note, while *C. intermedius* has not been measured for urine concentration, C. *penicillatus* is its sister taxon and is found in the same environment. B. Phylogenetic relationships of target species coded by habitat type (desert in orange, non-desert in green). Divergence time estimates from TimeTree.org.

We sequenced kidney mRNA from these pairs of taxa and assembled and annotated *de novo* transcriptomes for 3 desert-mesic species pairs spanning ∼70 million years of evolution. Assembled transcriptomes were used to analyze rates of evolution in single copy orthologs to identify genes putatively under selection across desert lineages. We also performed mRNA-sequencing on multiple individuals within each species to study gene expression divergence between desert and non-desert species. Global patterns of gene expression recapitulated the phylogeny of these six species. However, we also discovered a significantly greater number of convergent changes in gene expression than expected by chance between desert and non-desert species. In contrast, convergent changes in amino acid sequence were identified at a smaller proportion of genes. Overall, we identified genes with shared patterns of molecular evolution associated with habitat type.

## Materials and Methods

### Sample collection

Five adult male mice for each species, with the exception of *H. desmarestianus*, for which only four samples could be obtained, were included in this study. *C. intermedius, Z. princeps*, and *M. musculus* were caught by N. Bittner using Sherman live traps set over-night following the guidelines of the American Society of Mammalogists (Sikes and Gannon 2011) and an ACUC protocol approved by UC Berkeley (AUP-2016-03-8536). Animals were given apple after capture to avoid dehydration for the short period of time they were in traps. Mice were euthanized by cervical dislocation, and kidney and liver were removed and preserved in RNAlater. *C. intermedius* were trapped near Tucson, AZ, USA, *Z. princeps* were trapped at Sagehen Creek Field Station near Truckee, CA, USA, and *M. musculus* were trapped near Berkeley, CA, USA. *H. desmarestianus* were collected in Chiapas, Mexico by Beatriz Jimenez, and *N. alexis* were collected by Kevin Rowe in Northern Territory, Australia. Mice collected by N. Bittner were prepared as museum specimens (skins and skulls) and deposited in the collections of the UC Berkeley Museum of Vertebrate Zoology (MVZ). Animals collected by K. Rowe were prepared as museum specimens and deposited at Museums Victoria. The collecting localities, collector’s numbers, and museum catalog numbers for each specimen are provided for all wild-caught animals in Table S1. Samples from *Jaculus jaculus* were provided by Kim Cooper at UC San Diego from an outbred lab colony. Despite the fact that the *Jaculus* were from a laboratory colony while all other animals were wild caught, patterns of gene expression among all individuals reflected the phylogeny of these taxa, suggesting that the laboratory environment for *Jaculus* did not obscure overall expression patterns (see Results).

### mRNA library preparation and sequencing

To target loci underlying adaptation to xeric environments, we focused on genes expressed in the kidney. RNA was extracted from kidney preserved in RNAlater using the MoBio Laboratories Powerlyzer Ultraclean Tissue & Cells RNA Isolation Kit. Remaining DNA was removed with DNAse-1 followed by a Zymo RNA Clean and Concentrator column clean-up. Due to the poor quality of some samples (RIN scores below 5), a ribosomal RNA depletion step was performed with a KAPA Riboerase Kit before libraries were prepared with the KAPA HyperPrep Kit. Libraries were pooled and sequenced across two lanes of 150 bp PE NovaSeq (one lane of S1 and one of SP) at the Vincent J. Coates Genomics Sequencing Center at UC Berkeley. One library from each species (except *Mus musculus*; see below) was sequenced at greater depth for transcriptome assembly; these were sequenced to a target of 100M read pairs while the remaining 24 libraries, intended for expression analysis, were sequenced to a target of 20M read pairs (see File S1).

### Transcriptome assembly

For each of the five 100M-read-pair libraries, reads were examined for quality metrics with FastQC (https://www.bioinformatics.babraham.ac.uk/projects/fastqc/) and then corrected by removing erroneous k-mers using rCorrector (Song and Florea 2015). Adapters and poor quality sequence were trimmed using Trim Galore! (https://www.bioinformatics.babraham.ac.uk/projects/trim_galore/). Since FastQC revealed a large quantity of duplicates within the sequenced libraries, which is likely in part due to rRNA contamination, we chose to remove all reads that mapped to known rodent rRNA from NCBI using bowtie2 (Langmead and Salzberg 2012). We ran Trinity v2.1.1 (Grabherr et al. 2011) to generate a transcriptome assembly for each species. Because transcriptome-depth (i.e. 100M read pairs) sequencing was not done for *Mus musculus*, reads from all five individuals (approximately equal to the sequencing depth for transcriptome individuals) were combined to assemble the transcriptome of a local individual as above (Table S2). To remove redundant transcripts from the Trinity assembly, transcripts with equal to or greater than 95% sequence identity were clustered with cd-hit-est (settings: -c 0.95 -n 8) (Li and Godzik 2006) to create representative transcripts before use in downstream analysis (Table S2). This was done to collapse transcript isoforms as well as to remove transcripts created by assembly errors (chimeras, duplicates, misassembled transcripts and the like). Transrate (Smith-Unna et al. 2016) was used to calculate assembly statistics. To assess assembly completeness, we used Benchmarking Universal Single-Copy Orthologs (BUSCO) (Seppey et al. 2019) to look for the 6,192 orthologs found in the Euarchontoglires odb9 database and thus expected to exist in the taxa studied here.

### Transcriptome annotation and ortholog detection

To identify coding regions within our assembled transcripts for downstream analyses, we utilized TransDecoder v. 5.5.0 (http://transdecoder.sourceforge.net). We identified the longest open reading frame (ORF) and searched for matches to both the Pfam protein domain database (Bateman et al. 2004) and mouse specific SwissProt database (Bairoch and Apweiler 2000) to retain ORFs based on homology. Since high quality gene annotations were available for *M. musculus*, we used the curated RefSeq protein database for this species. Orthologous gene groups across all six taxa were identified using OrthoFinder v 2.3.3 (setting: -S diamond) (Emms and Kelly 2015). To minimize the number of alternate isoforms used in the analysis, we only used the longest ORF identified per gene.

### mRNA read mapping

Raw reads from all libraries were examined for quality with FastQC. Adapters and poor quality sequence were trimmed using Trimmomatic v0.36 (Bolger et al. 2014). The five libraries that were generated for transcriptome assembly were subsampled to the average read number of the libraries generated for expression (27,787,405 reads). Reads were mapped to transcriptomes generated for each species with Salmon v 0.14.1 (Patro et al. 2017). To compare across genera, transcripts were annotated using BLASTn to the Refseq cdna database for *Mus musculus*. Read counts were summed across transcripts for each annotated gene.

### Quantification of gene expression and identification of convergent differential expression

DESeq2 (Love et al. 2014) was used to normalize for differences in library size and to call differential expression between species within each family and across all samples. As transcripts between species can differ in length, a length correction was applied. Reads were subsequently transformed with a variance stabilizing transformation for principal component analysis.

We used DESeq2 to identify convergent changes in gene expression between desert and non-desert species across all three families using an approach similar to that used by Parker et al. (2019). In particular, we fit a generalized linear model for gene expression as a function of habitat (desert vs. non-desert), family (species-pair), and their interaction. Genes were classified as convergently differentially expressed in cases where there was a significant effect of habitat (desert vs. non-desert, FDR<0.01) but no interaction effect of species-pair by habitat (FDR > 0.05). This analysis was restricted to genes with greater than an average of 20 reads per sample for each species, resulting in a total of 8,174 genes. P-values were adjusted for multiple testing using a Benjamini & Hochberg (Benjamini and Hochberg 1995) correction. Differential expression within each species pair was identified using pairwise contrasts. For pairwise contrasts, genes with a mean of fewer than 10 reads per sample were removed from the analysis.

Permutation tests were used to assess whether more genes showed convergent shifts by habitat type than expected by chance, as described in Parker et al. (2019). For each gene, read counts were randomly assigned to habitat within each species pair. All biological replicates (i.e. all five individuals) in each species were assigned to the same habitat. This process was used to create 10,000 permuted datasets. The number of convergently differentially expressed genes in these datasets were compared to that of the observed dataset

### Estimating rates of molecular evolution and identification of genes under positive selection

Using the single copy ortholog groups generated by OrthoFinder for all six species, we aligned these using MAFFT through Guidance2 (Sela et al. 2015), which provided alignment quality scores for all 1,855 genes and removed those for which the alignment quality score was poor (mean column score <0.8). Alignments for which quality scores were poor were removed from subsequent analyses, resulting in a set of 1,474 genes with aligned protein coding alignments for subsequent analyses.

We used a maximum likelihood approach in a phylogenetic context by implementing the codeml package in PAML (Yang 1997) to identify genes in desert lineages with evidence of selection. We performed three analyses using the 1,474 single copy orthologs with high quality alignments present in all species. First, we defined the three desert species together as “foreground” lineages and compared these to the three non-desert species as “background” lineages using a foreground-background branch analysis implemented in PAML (Yang 1998; Yang 2007). This analysis estimates ω or dN/dS (the rate of nonsynonymous substitutions per nonsynonymous site divided by the rate of synonymous substitutions per synonymous site) and compares branches of interest (e.g., “foreground” branches) to the other “background” branches. Elevated rates of dN/dS compared with a null model are considered evidence for selection. This analysis was intended to identify genes underlying desert adaptation common to all three species. Second, we performed three separate foreground-background branch analyses, in which each desert species by itself was compared to the other five species. This analysis was intended to identify species-specific adaptations. Third, we performed a branch site model which allows for ω to vary both across sites in a gene and across branches on the tree.

### Enrichment analyses

For gene sets of interest, GO category enrichment tests were performed with GOrilla (Eden et al. 2009) to test a foreground gene set of interest against a background set of all other genes included in the analysis. Phenotype enrichment tests were performed with modPhea (Weng and Liao 2017) using the same framework.

## Results

### Sequencing, assembly, and annotation

We generated on average ∼123 million reads per sample for the assembly of *de-novo* kidney transcriptomes in each species. For *Mus musculus*, five smaller libraries were concatenated for assembly. After read correction, quality filtering, and adapter trimming, each library had an average of ∼103 million reads which were used for the assembly. Each assembly contained 965,227 transcripts on average. We reduced the number of redundant transcripts in the assembly to improve accuracy of downstream analyses by clustering similar transcripts together using CD-HIT-EST. This decreased the number of transcripts by ∼20% per sample to an average of 793,887 transcripts (Table S2). We used BUSCO to check assembly completeness to determine how many of the 6,192 orthologs found in the Euarchontoglires odb9 were present in our assembled transcriptomes. The six assemblies ranged in completeness from 80 - 87% (Figure S1). This level of completeness reflects a single tissue (kidney) taken at one developmental time point. After ORF prediction, we annotated each transcript to known *M. musculus* proteins. We were able to assign transcripts to 395,029 putative ortholog groups.

### Global gene expression reflects phylogenetic relationships and habitat type

To identify patterns of differential gene expression, we sequenced kidney mRNA from additional individuals in each of the six species for an average of ∼27 million reads per individual. We retrieved 13,305 genes in *C. intermedius*, 11,749 genes in *H. desmarestianus*, 14,891 genes in *J. jaculus*, 14,380 genes in *Z. princeps*, 18,622 genes in *N. alexis* and 19,913 genes in *M. musculus* for which we were able to quantify expression levels. These genes were annotated using *M. musculus* transcripts so as expected, the number of genes we were able to annotate in more divergent species is more limited.

Gene expression profiles largely recapitulated the known phylogenetic relationships of these six species (Figure 2A). Individuals within each species form well-defined clusters (with the exception of a single *Heteromys* individual), and the different genera within each family share expression profiles that are more similar to each other than they are to genera in different families. Further, Muridae and Dipodidae are more similar to each other in expression profiles than either is to Heteromyidae, reflecting the known evolutionary relationships of these families. Thus, the overall expression patterns reflect evolutionary history more than habitat type. These patterns are also seen in a principal component analysis (PCA) based on expression level co-variance (Figure 2B), where PC1 (accounting for 33% of the variance) largely reflects phylogeny. Despite the overall phylogenetic pattern of gene expression, consistent differences in expression were seen between desert and non-desert species within each family. In particular, PC4 captures this variation, separating desert from non-desert taxa (explaining 11% of the variation) (Figure 2C).

**Figure 2.**
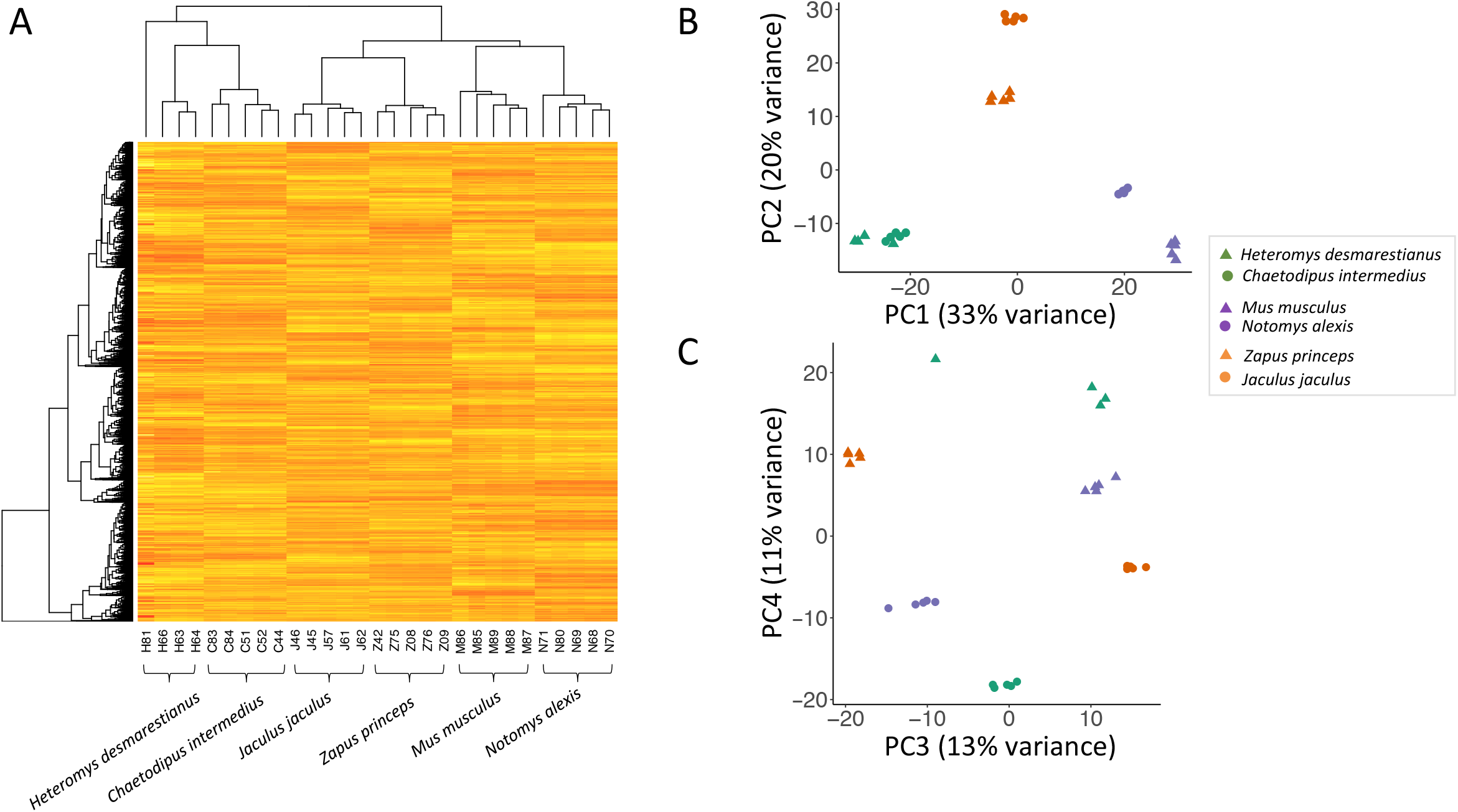
Expression level variation differentiates species and habitat type. A) Heat map showing relationships among samples based on gene expression clustering. With the exception of one sample (H81), expression patterns reflect phylogenetic relationships (see Figure 1). B) Principal components (PC1 and PC2) for the expression data. PC1 explains 33% of the variance and reflects the phylogenetic relationships of the species. PC2 explains 20% of the variance. C) Principal components (PC3 and PC4) for the expression data. PC4 explains 11% of the variance and differentiates samples by habitat type.

### Convergent differential expression in desert rodent kidneys

We quantified differential expression (DE) between desert and non-desert species within each family. In pairwise contrasts between desert and non-desert species in Heteromyidae, Dipodidae, and Muridae, we identified >4,000 genes in each comparison with evidence of significant DE (Table S3, FDR<0.01). Individual pairwise comparisons between desert and non-desert species found uniquely in each of the three families (to the exclusion of the two others) were associated with several GO categories, including cellular metabolic processes and nitrogen metabolic processes (Table S4). We identified a total of 654 genes that showed significant differential expression in all three species pairs (Figure S2), with 145 of these genes showing shifts in the same direction in each comparison.

To identify convergent shifts in gene expression associated with desert-living, we also modeled gene expression as a function of species pair (i.e., family), habitat, and their interaction. Convergent changes were identified as those for which there was a significant effect of habitat (FDR<0.01), but no interaction between species pair and habitat (FDR>0.05) (see Methods; Parker et al. 2019). We identified 702 genes with shared shifts in desert rodents relative to the mesic comparison (Figure 3A). This set includes all of the 145 genes identified above in pairwise tests. Thus, 8.6% (702/8,174) of genes showed convergent shifts in expression in desert rodents compared to their non-desert relatives.

**Figure 3.**
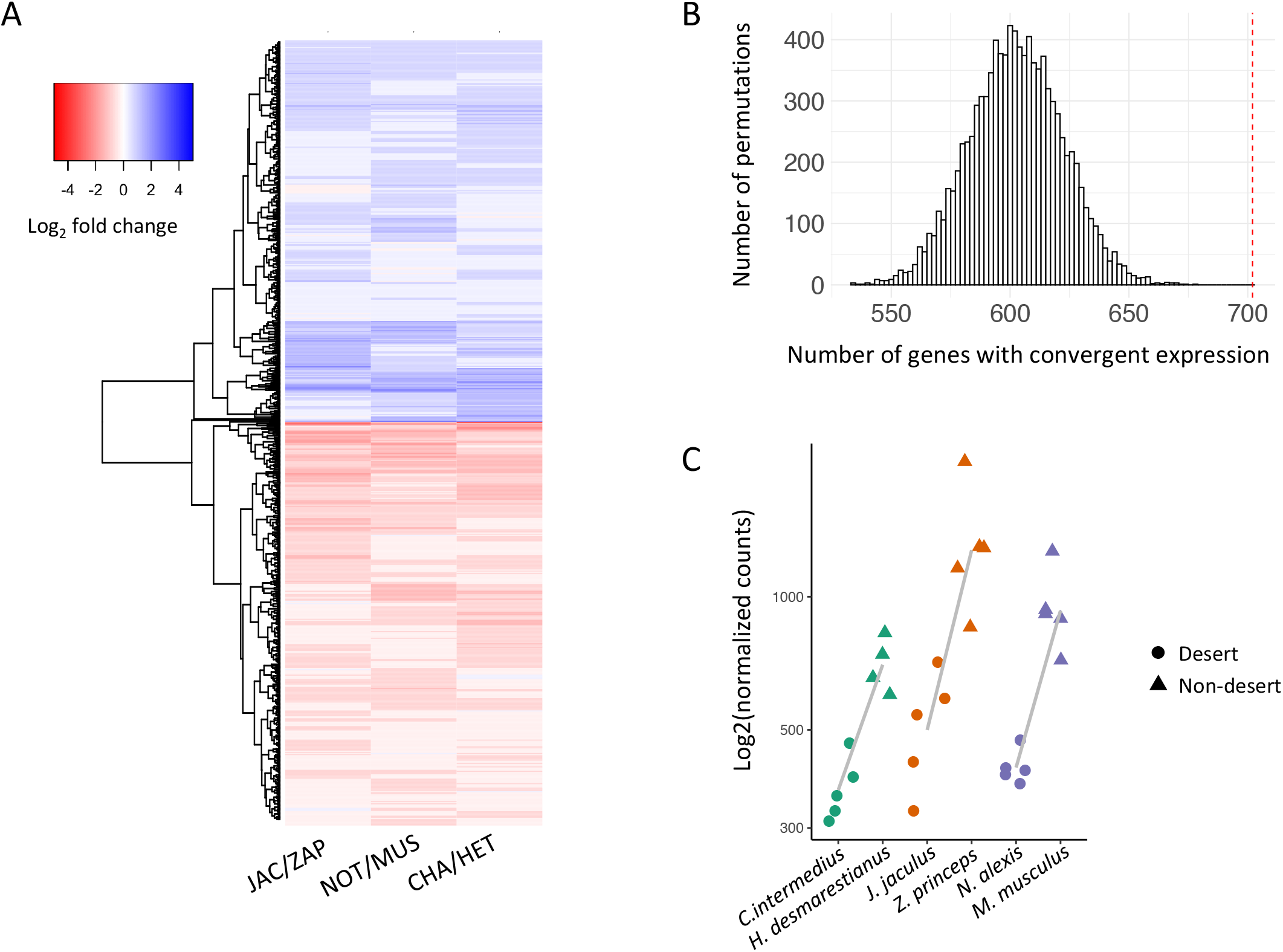
A) Heatmap of genes with evidence of convergent gene expression patterns. Each row is a gene. Each of the three columns shows the mean expression value among all desert individuals compared to the mean expression value of all non-desert individuals for each family. B) Number of genes expected by chance to show convergent expression after 10,000 permutations. Observed number of genes (red line) is greater than the distribution expected by chance (*p* < 0.0001). C) Expression values for *Aqp11*, a gene showing convergent gene expression. In all comparisons, desert species show lower expression levels compared to their non-desert relative. Grey dotted lines are drawn between the mean of normalized expression for each desert-mesic pair. Points are jittered for clarity.

Shared shifts in gene expression can be a consequence of selection in response to shared environmental pressures or stochastic processes. To ask if the observed number of genes with convergent differential expression was more than expected by chance, we performed a permutation test in which we took each gene and randomly switched habitat assignment within species pairs, while always maintaining the same label for all biological replicates within a species, to create 10,000 permuted data sets (Figure 3B, see Methods). Permuted datasets never identified more convergent genes than the observed set of convergent genes, suggesting an enrichment of convergent differential expression associated with habitat type.

Fold changes between individual desert-mesic pairs were often modest in one or more contrasts between species pairs (Figure S3); only 208 genes with shared expression shifts showed an average of greater >0.5 log2 fold change difference between each desert-mesic species pair. The number of genes showing higher expression in desert rodents compared to non-desert relatives (335 genes, shown in blue in Figure 3A) was slightly fewer than the number of genes showing lower expression in desert rodents compared to non-desert relatives (367 genes, shown in red in Figure 3A). Additionally, across all genes, fold changes between individual desert-mesic species were found to be significantly correlated in 2 of the 3 comparisons of species pairs (Spearman’s rank correlation rho, *C. intermedius/H. desmarestianus* vs. *J. jaculus/Z. princeps, p=*0.0062, *rho=*0.03; *C. intermedius/H. desmarestianus* vs. *N. alexis/M. musculus, p*< 2.2e-16, *rho*=0.10; *N. alexis/M. musculus* vs. *J. jaculus/Z. Princeps, p=*0.12, *rho=*-0.017).

To identify genes and pathways of interest, we divided the set of convergently expressed genes into those that are upregulated with respect to the desert taxa in all comparisons and those that are downregulated with respect to the desert taxa in all comparisons and performed phenotype and GO term enrichment tests on these (see methods). Genes convergently upregulated across desert rodents were enriched for several GO terms related to gene regulation, including regulation of RNA metabolic process (*q*=2.55 × 10^−5^), regulation of gene expression (*q*=1.34E-5), and regulation of RNA biosynthetic process (3.87 × 10 ^5^). Genes downregulated in desert rodents were enriched for GO terms related to metabolic processes, including metabolic process (*q*=1.56 × 10 ^3^), organic substance metabolic process (*q*=3.93 × 10^−3^), and cellular metabolic process (3.54 × 10^−3^). Genes with evidence for convergent differential expression included genes with mouse mutant phenotypes related to kidney development and physiology or homeostasis (Table S5). For example, Aquaporin 11 (*Aqp11*) is expressed at a lower level in all desert species compared to non-desert species in all three comparisons (Figure 3C). This gene is part of a family of genes encoding membrane-integrated channels responsible for water transfer across membranes throughout the body. Aquaporins have been repeatedly implicated in studies of desert adaptation across rodents (Marra et al. 2012; Marra et al. 2014; Pannabecker 2015; Giorello et al. 2018). Mouse knockouts have demonstrated that *Aqp11* is necessary for proximal tubular function and the formation of healthy kidneys (Morishita et al. 2005; Tchekneva et al. 2008). In addition, *Aqp11* plays a role in salivary gland development (Larsen et al. 2010). This set also includes genes associated with human phenotypes related to kidney and renal diseases (Table S6); for example, mutations in the gene *col4a5*, which is downregulated in desert species, have been associated with Alport syndrome, a disease characterized by kidney inflammation (Köhler et al. 2019).

### Genes under selection in desert lineages

Next, we tested for evidence of selection on protein coding sequences using well aligned one-to-one orthologs found in all desert-mesic species pairs (1474 genes). We searched for genes showing signatures of selection using a model that compares the rate of nonsynonymous substitutions with the rate of synonymous substitutions in a phylogenetic context, ω or dN/dS (Yang, 1998, 2007). We performed three analyses using the 1,474 single copy orthologs with high quality alignments present in all species. When testing for evidence of selection on desert species compared to non-desert species, we uncovered 39 genes (39/1474= 2.6%) for which ω was significantly higher in the three “foreground” desert lineages compared with the three “background” non-desert lineages (Table S7, FDR<0.1). This group is enriched for phenotypes related to multiple aspects of the immune response as well as to hearing/vestibular/ear phenotypes and other aspects of osteology (Table S8). Immune genes are some of the fastest evolving genes in the genome and are disproportionately found to be under selection in many studies (Hurst and Smith 1999; Schlenke and Begun 2003; Nielsen et al. 2005). One gene of particular interest, unrelated to immunity, is FAT atypical cadherin 4 (*FAT4*) (*q* = 0.018). *FAT4* has been implicated in human kidney diseases (Alders et al. 2014) and is involved in normal kidney development through modulating the RET signaling pathway in mouse models (Mao et al. 2015; Zhang et al. 2019). *FAT4* homozygous knockout mice have smaller kidneys with the presence of cysts in renal tubules when compared with wild type mice and they die within a few hours of birth (Saburi et al. 2008). These phenotypes in laboratory mice make this an interesting candidate gene for future studies in desert rodents. We found three genes that showed evidence of positive selection (when the three desert species were treated together as foreground lineages) and also showed convergent shifts in gene expression (Rows 1-3 in Table S9), however they are not known to be associated with phenotypes of interest. This amount of overlap is no more than expected by chance (hypergeometric test, *p*=0.64).

We then tested whether ω was significantly higher in each of the three “foreground” desert lineages individually compared with the five remaining taxa. We identified 23 genes in *C. intermedius*, 19 in *J. jaculus*, and 18 in *N. alexis* where ω was significantly elevated (at FDR <0.1)(Table S10). These genes are candidates for lineage-specific adaptations. In *C. intermedius*, enriched phenotypes were related to immunity and morphological traits including kidney size, while in *J. jaculus* and *N. alexis*, enriched phenotype terms were related to behavioral and electrophysiological traits (Table S11). In the *Chaetodipus* comparison, *Dusp4* is of some interest as it has been associated with aberrant circulating solute levels in mouse models. Deletion of this gene has been associated with increased excreted protein and altered kidney structure in diabetic mice (Denhez et al. 2019). It is also convergently differentially expressed. Overall, the amount of overlap (hypergeometric test, *p* > 0.06 in all comparisons) between any of these lists and differentially expressed genes between lineage pairs is no more than expected by chance (Table S9).

In the third analysis, we employed a branch-site model to identify genes in which specific codons may be under positive selection. In this approach, genes for which specific codons have a ω > 1 in the “foreground” branch (defined to include all three desert species) compared with the “background” branch are identified. Seven genes were identified (Table S12) with codons under selection in all three desert lineages, including *Coro2b*, a gene implicated in abnormal renal glomerulus morphology (Schwarz et al. 2019) and urine protein level (Rogg et al. 2017) and *Bloc1s4*, which is implicated in abnormal renal physiology (Gwynn et al. 2000). Again, there was no significant overlap with the genes identified in the differential expression analysis (*p*=0.47; Table S9).

## Discussion

The molecular basis of convergent evolution has been well studied for a number of simple traits, but has been less studied for complex traits. Even fewer studies have compared convergence in both gene expression and protein evolution for complex traits. Here, we studied convergence in gene expression and amino acid sequence in three species of desert rodents and their non-desert relatives, from across the rodent tree, representing ∼70 million years of evolution.

Despite the long evolutionary timeframe and the fact that most expression evolution tracked phylogeny (Figure 2), we identified a surprising number of genes (702/8174=8.6%) that showed convergent shifts in gene expression (Figure 3). This number is more than expected by chance, and this result is highly significant (Figure 3B). We note, however, that the number of genes showing convergent expression does not reflect the number of causative changes (i.e. mutational events in evolution), since many of these convergent changes in expression might reflect downstream consequences of a smaller number of changes at upstream regulators that govern networks of co-regulated genes. Nonetheless, the large number of convergent changes in expression suggests that a measureable amount of desert adaptation is mediated by a large set of shared changes in gene regulation, whether at the level of individual genes or through sets of co-regulated genes.

In addition to these shared changes in gene expression, we identified a large number of species-specific changes in gene expression in each species pair. Perhaps not surprising given the long evolutionary timescales and complexity of osmoregulatory function, much of the evolutionary response appears to be specific to individual lineages.

In contrast to the fairly long evolutionary timescales in the present study, we recently documented differences in kidney gene expression between desert and non-desert populations of *Mus musculus* separated by only a few hundred generations of evolution (Bittner et al. 2021). In that study, we identified 3,935 differentially expressed genes of which 99 were found to be convergent across all three desert lineages in the present study. The lack of significant overlap (hypergeometric test, *p*=0.99) suggests that over long evolutionary timescales, adaptive responses to xeric conditions may be quite different from the evolved changes in gene expression over short evolutionary timescales.

In contrast to the number of convergent changes in gene expression, we observed few genes that showed evidence of positive selection on amino acid sequences (39/1474= 2.6%) among desert species. These proportions are not directly comparable since the methods used to detect convergence and positive selection are quite different. Nonetheless, our analyses suggest that the phenotypic convergence seen in urine concentration is reflected at the molecular level more in patterns of gene regulation than in patterns of protein evolution.

Although expression evolution and amino acid sequence evolution have been found to be correlated in some cases (Nuzhdin et al. 2004; Lemos et al. 2005), we did not find significant overlap in the number of genes showing convergent gene expression and convergent amino acid sequence evolution. The small amount of overlap might reflect differences in the selection pressures on these two classes of changes. For example, *cis*-regulatory changes in gene expression are often controlled in a tissue-specific and developmental-stage-specific manner, and as such are expected to be less pleiotropic and thus less constrained in evolution (e.g. Wray 2007). Protein-coding changes, on the other hand, affect all tissues and developmental stages in which the protein is expressed and thus may be more pleiotropic and consequently more constrained. The small amount of overlap might also reflect both statistical and methodological limitations of our study. First, the analytic methods used to detect convergent expression changes and convergent amino acid changes are quite distinct and likely have different false-negative and false-positive rates. Second, we studied gene expression in adults, yet gene expression varies considerably during kidney development (Schwab et al. 2003) and early expression is undoubtedly important in establishing morphological differences between desert and non-desert kidneys. Third, kidneys have a heterogenous cellular composition, and changes in cellular composition between species are likely to affect measures of gene expression in bulk preparations. Future studies of gene expression in single cell preparations at early developmental time points might uncover additional signals of adaptation to desert conditions. It would also be worthwhile to study expression changes in both males and females since the demands of water balance may be especially acute in lactating females.

Despite these caveats, we identified a number of potential candidate genes associated with desert adaptation, including some that showed both convergent gene expression and convergent amino acid sequence evolution. The target available to selection in a trait as complex as desert adaptation is likely large and constrained along each lineage to a different degree by other aspects of the organism’s morphology and physiology. Nonetheless, an interesting outcome of our analysis is that a number of the genes and pathways identified here are similar to those identified in other studies of rodent and mammalian desert adaptation (Marra et al. 2012; Marra et al. 2014; Wu et al. 2014; MacManes 2017; Giorello et al. 2018; Tigano et al. 2020). It is clear that gene families such as aquaporins, which are responsible for facilitating water transport across membranes, and solute carriers, may play a role in mitigating water loss across multiple systems and therefore underlie convergent evolution at the genetic level to desert environments. Altogether, our results demonstrate the power of studying convergent evolution at multiple levels by integrating scans for convergent evolution on both the amino acid and gene expression level to identify genes and pathways of interest. Our results suggest that changes in gene regulation may play an essential role in how the kidneys of species separated by 70 million years of evolution have convergently evolved ultra-efficient osmoregulatory capacity to contend with the stressors of harsh desert environments.

## Acknowledgements

We thank members of the Nachman Lab for valuable comments and discussions. We thank Dr. Kim Cooper, Dr. Kevin Rowe, and Dr. Beatriz Otero Jiménez for generously sharing their samples as well as Dr. Andy Gloss, Dr. Shea Lambert, Dr. Aaron Ragsdale, Dr. Taichi Suzuki, Ned McAllister and the Oakland Zoo, and Kim Hemmer and Golden Gate Fields for their help with field collections. We thank Dr. Jim Patton, Dr. Andy Gloss, Lydia Smith, and Jeremy Richardson for their technical expertise. This work was supported by a NSF Doctoral Dissertation Improvement Grant to NKJB (1601827), grants from the Wilhelm L.F. Martens Fund and the David and Marvalee Wake Fund of the Museum of Vertebrate Zoology, and an NIH grant to MWN (R01 GM127468).

## Data Accessibility Statement

Illumina sequencing data from this study is available is available through the NCBI Sequence Read Archive under accession PRJNA656179. Samples collected by NKJB are accessioned into the Museum of Vertebrate Zoology collection.

## Author Contributions

This study was designed by all authors. NKJB conducted the experiments, and NKJB and KLM analyzed the data. The paper was written by NKJB and edited by MWN and KLM.

## Supplemental material

**Figure S1.**
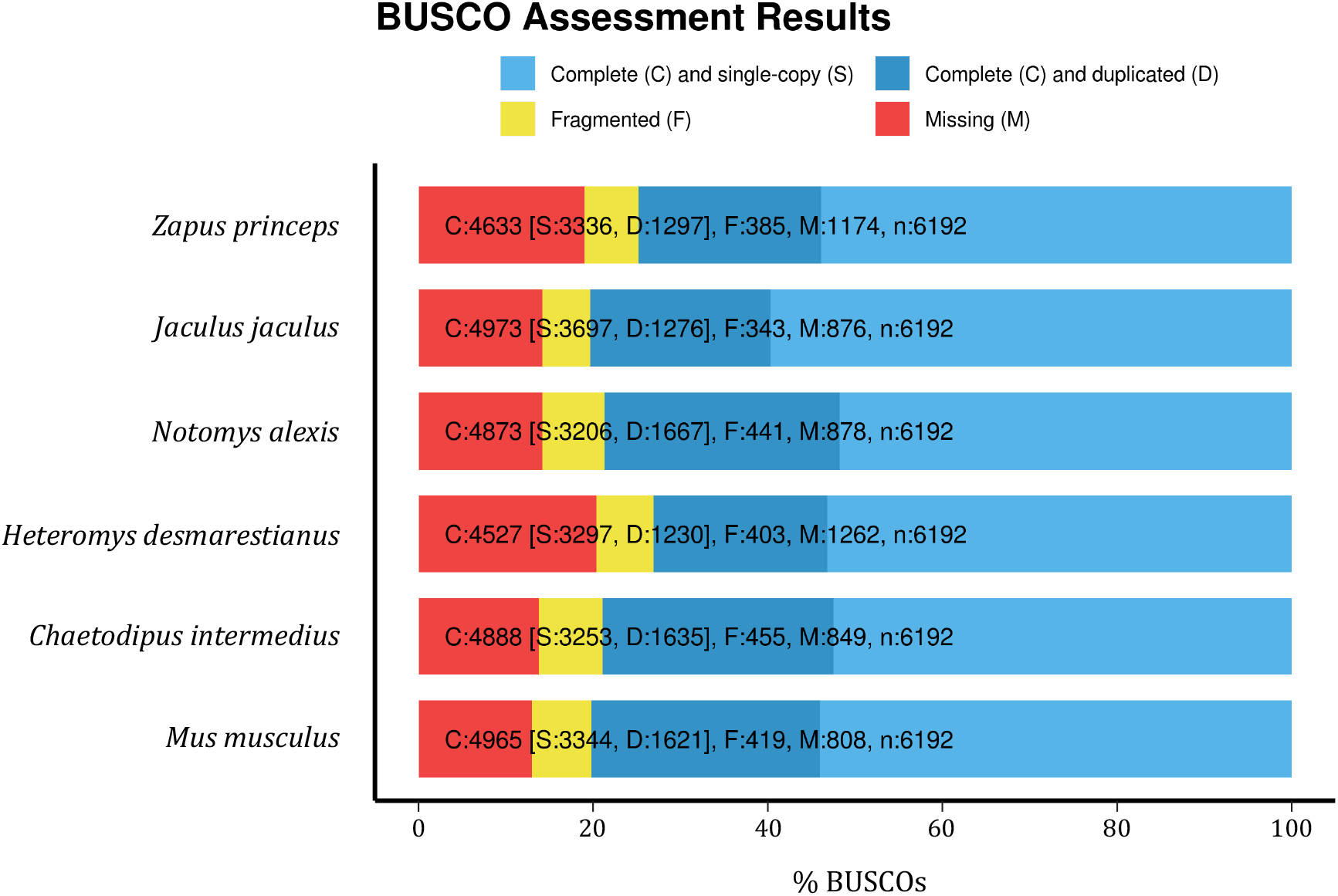
Benchmarking Using Single Copy Orthologs (BUSCO) score for each of the transcriptome assemblies

**Figure S2.**
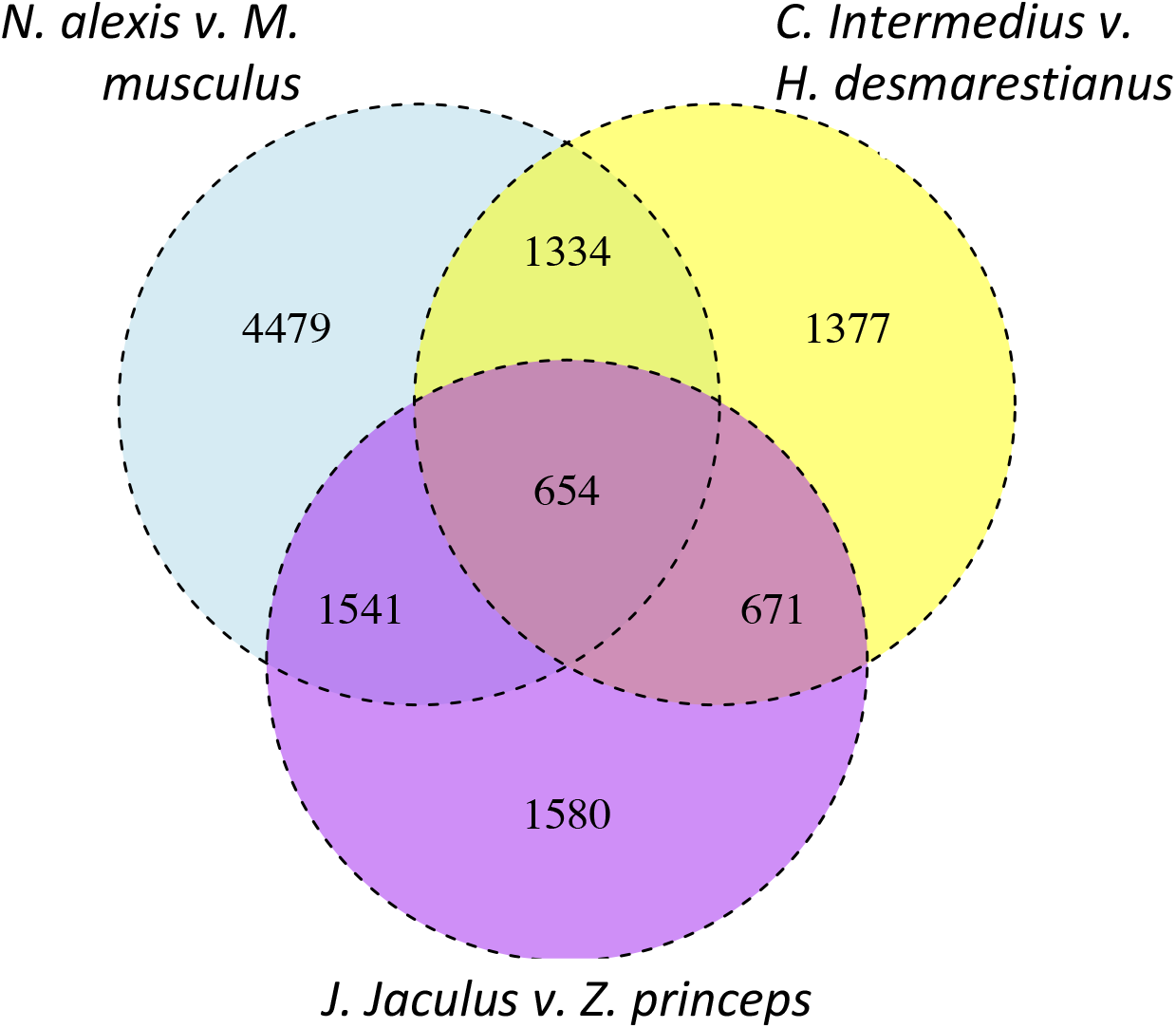
Genes that are differentially expressed between each desert and non-desert comparisons within each family and the overlaps among families.

**Figure S3.**
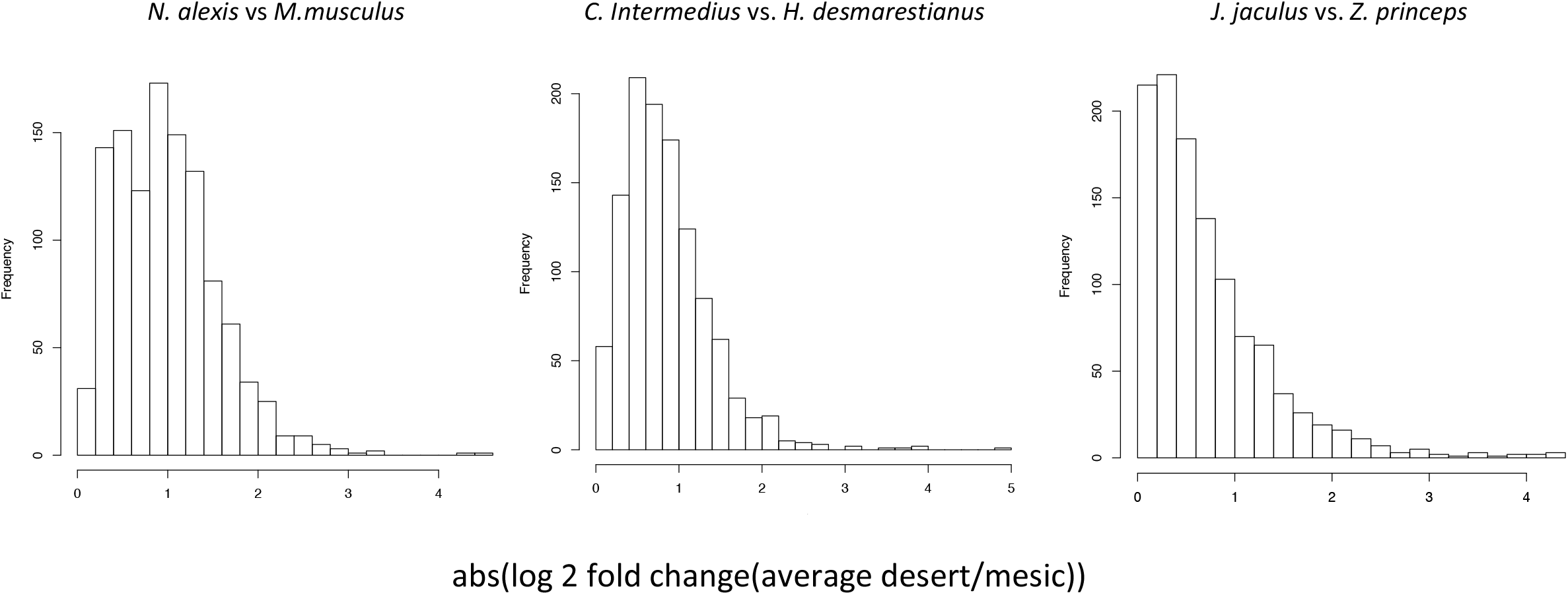
Magnitude of expression differences between each desert-mesic species pair

