## Supplemental Figures for "Convergent patterns of gene expression and protein evolution associated with adaptation to desert environments in rodents"

### Supplemental material

**Figure S1.** Benchmarking Using Single Copy Orthologs (BUSCO) score for each of the transcriptome assemblies

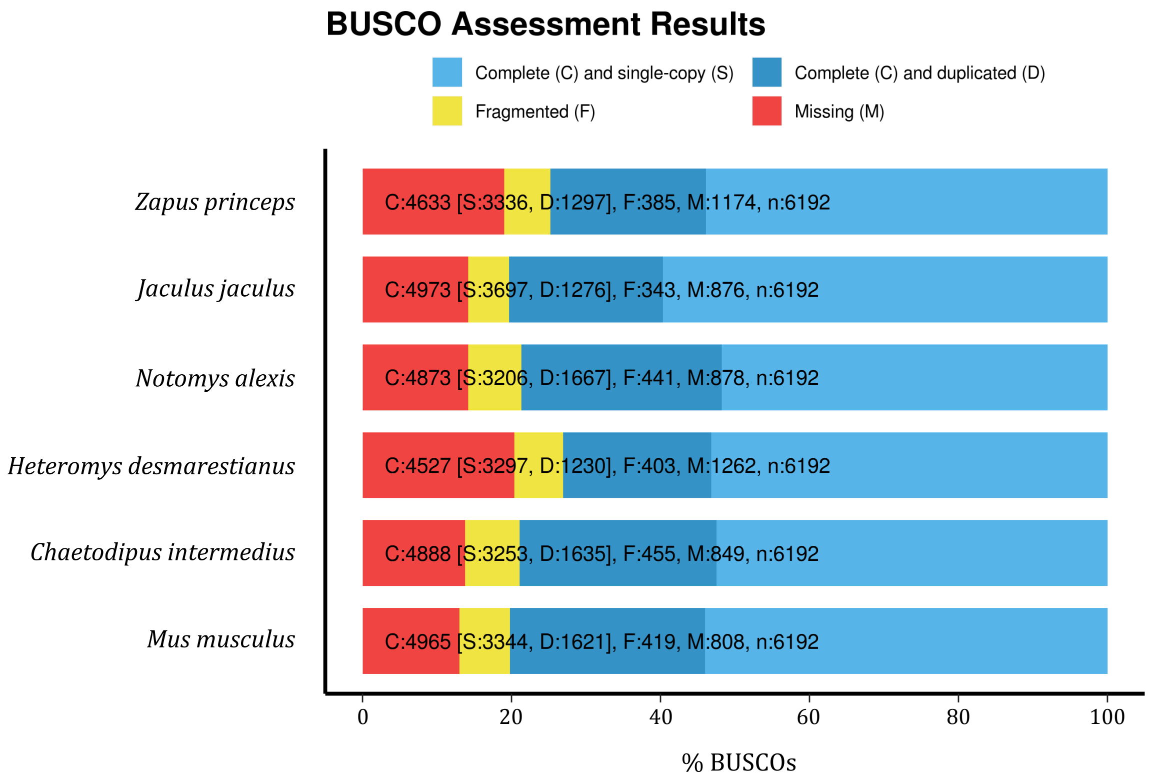

**Figure S2.** Genes that are differentially expressed between each desert and non-desert comparisons within each family and the overlaps among families.

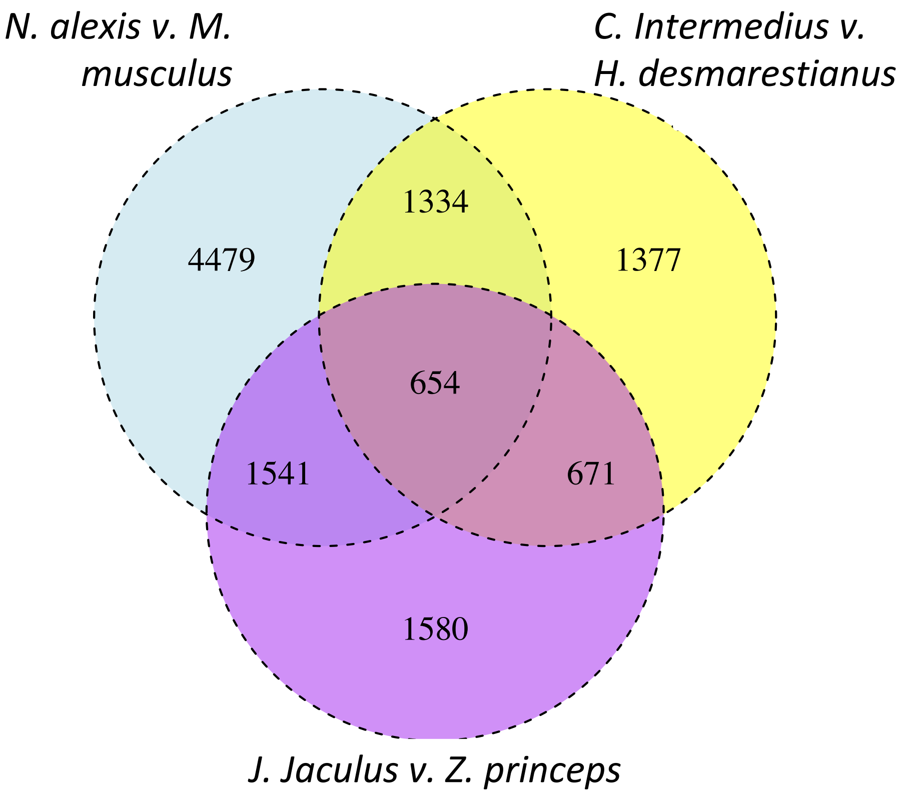

**Figure S3.** Magnitude of expression differences between each desert-mesic species pair

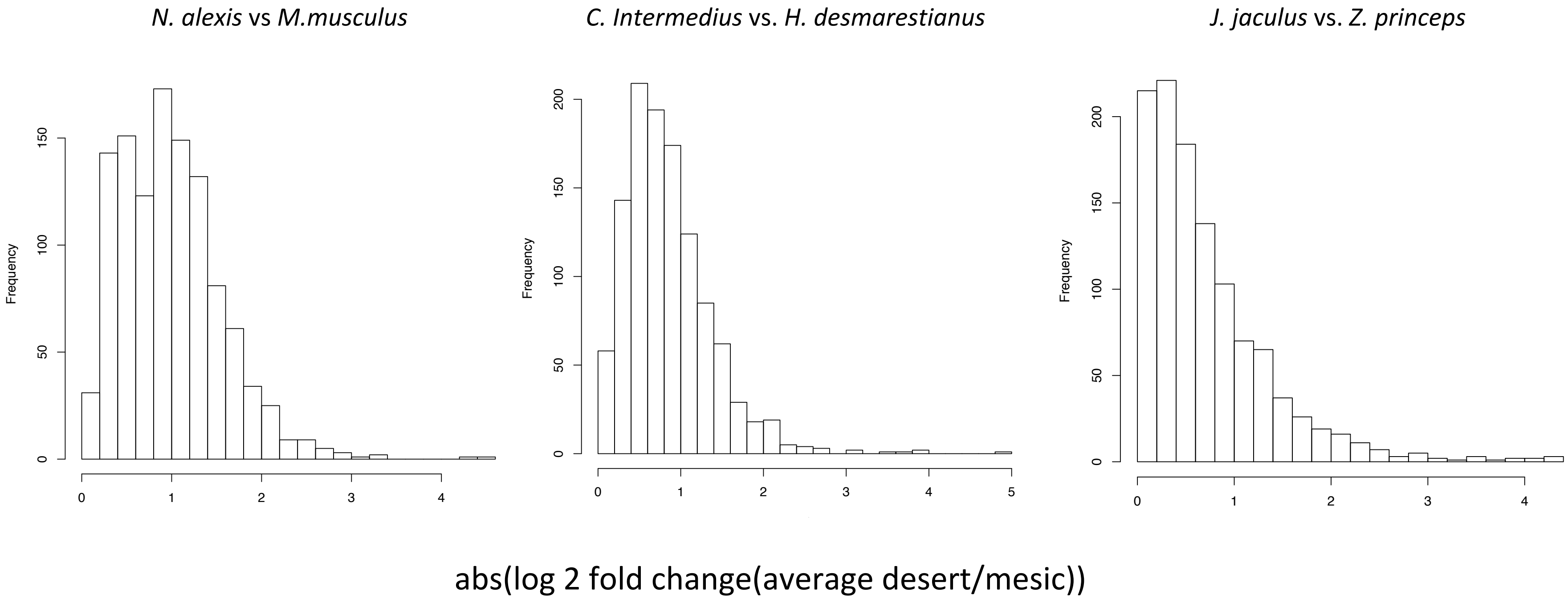
